## Supplementary information for "Maintenance of mitochondrial integrity in midbrain dopaminergic neurons governed by a conserved developmental transcription factor"

### Supplementary Figures

**Supplementary Figure 1**

**Supplementary Figure 2**

**Supplementary Figure 3**

**Supplementary Figure 4**

**Supplementary Figure 5**

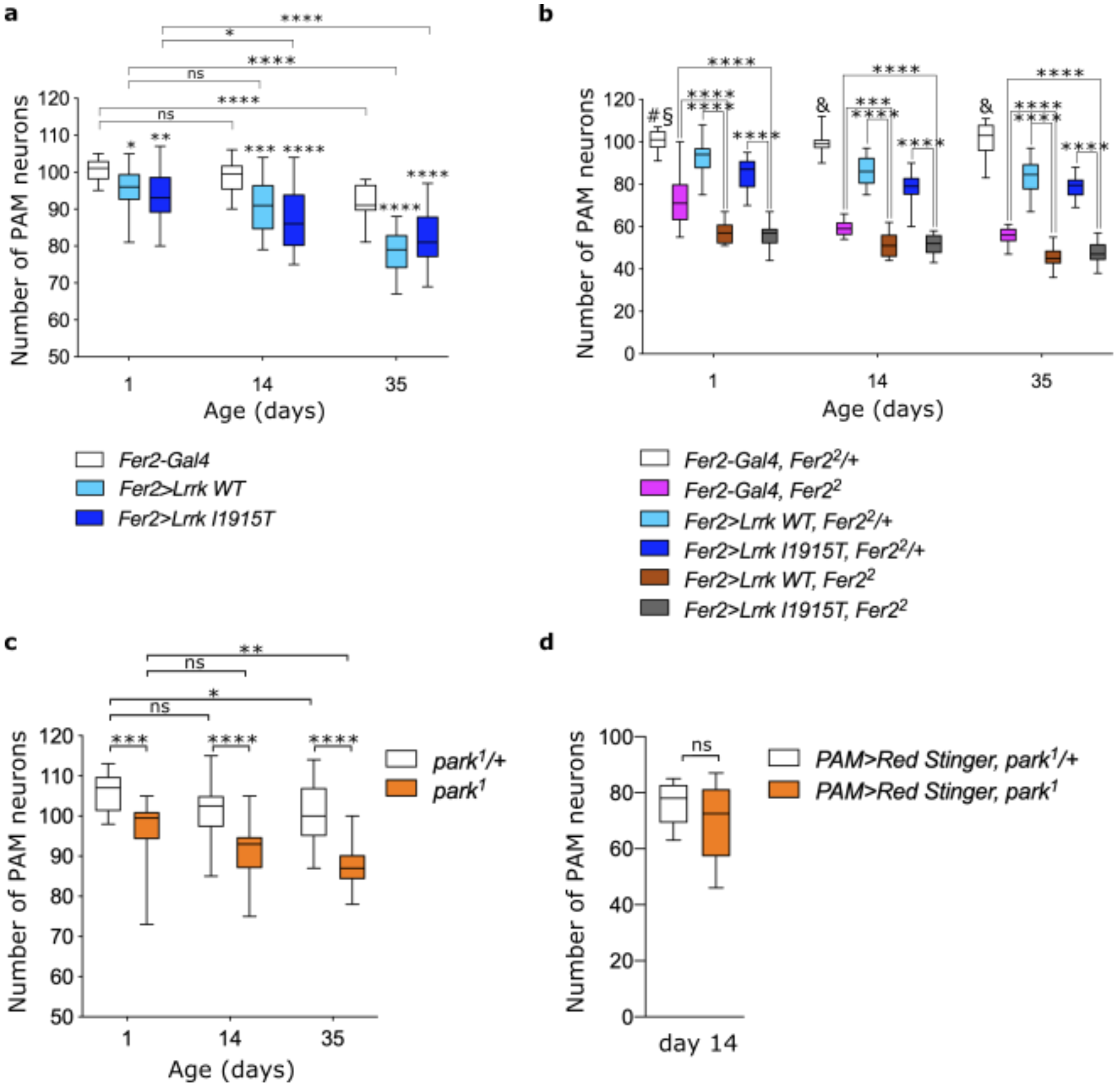


**Supplementary Figure 1. Quantification of PAM neuron counts *in LRRK2*-linked PD model flies and in *park*$^{1}$ mutants.**

**(a-c)** Quantification of the number of PAM neurons per hemisphere, as detected by anti-TH immunostaining for the indicated genotypes and ages. *Fer2>X* indicates that *UAS-transgene X* is expressed with *Fer2-GAL4*. n=14-20 hemispheres per group. Box boundaries are the 25th and 75th percentiles, the horizontal line across the box is the median, and the whiskers indicate the minimum and maximum values. One-way ANOVA followed by a Turkey's test for multiple group comparison., *p<0.05, **p<0.01, ***p<0.001, ****p<0.0001. ns, not significant. In (**a**), where brackets are not shown, flies overexpressing *Lrrk* and *Fer2-Gal4* flies (ctrl) of the same age are compared. In (**b**) at day 1, §p<0.05 when comparing *Fer2-Gal4, Fer2*$^{2}$/+ with *Fer2>Lrrk WT, Fer2*$^{2}$/+, and #p<0.0001 when comparing *Fer2-Gal4, Fer2*$^{2}$/+ with all the other genotypes. At day 14 and 35, &p<0.0001 when comparing *Fer2-Gal4, Fer2*$^{2}$/+ with all the other genotypes at the same age. **(c)** Quantification of TH-positive cells within the PAM cluster in homozygous and heterozygous *park*$^{1}$ mutants. *p<0.05, **p<0.01, ***p<0.001, ****p<0.0001. ns, not significant. n=14-20 hemispheres per group. One-way ANOVA followed by a Turkey's test for multiple group comparison. **(d)** Quantification of the number of PAM neurons labeled with RedStinger in *park*$^{1}$ mutants at day 14. The elements of the box plots are as in (a-c) and defined in the Method section. Mann-Whitney test found no difference between two genotypes.

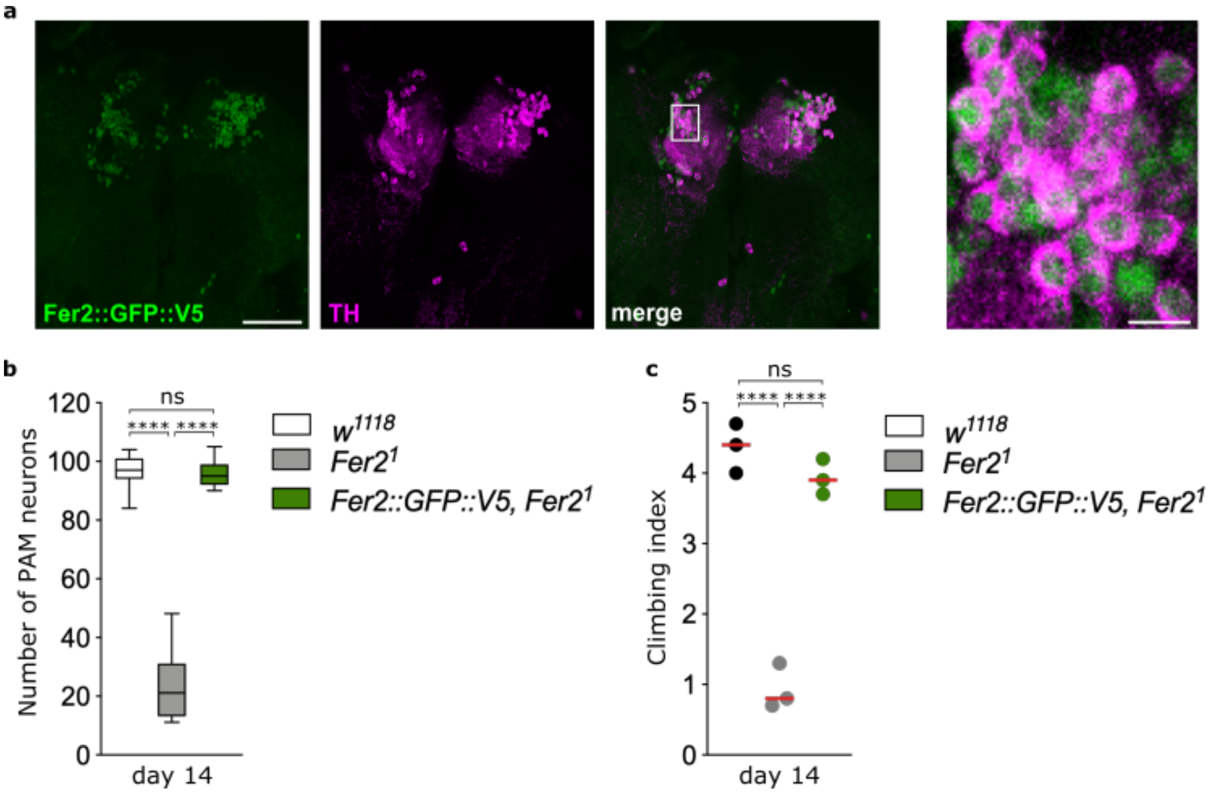


**Supplementary Figure 2. *Fer2::GFP::V5* transgene rescues PAM neuron loss and locomotor deficits in *Fer2*$^{1}$ mutants.**

**(a)** Brains of *Fer2::GFP::V5, Fer2*$^{1}$ flies at day 14 were stained with anti-GFP (green) and anti-TH (magenta) antibodies. The right panel presents a high-magnification image of the PAM neurons in the square from the left panel. Left panel scale bar, 50 µm. Right panel scale bar, 5 µm. **(b)** Quantification of the number of PAM neurons per hemisphere, as detected by anti-TH immunostaining in *w*$^{1118}$, *Fer2*$^{1}$ and *Fer2::GFP::V5, Fer2*$^{1}$ flies at day 14. n=14-20 hemispheres per group. One-way ANOVA followed by a Turkey's test for multiple group comparison, ****p<0.0001. ns, not significant. **(c)** Climbing index of *w*$^{1118}$, *Fer2*$^{1}$ and *Fer2::GFP::V5, Fer2*$^{1}$ flies at day 14. Three independent experiments. One-way ANOVA followed by a Turkey's test for multiple group comparison, ****p<0.0001. ns, not significant.

**a**

| Predicted TFs binding to PAM DEG | | | Interactions with *Fer2* direct targets |
|---|---|---|---|
| TF | p-value | TF binding site database | |
| *BEAF-32* | 0.0032 | Jaspar | - |
| *br* | 0.0009<br>0.0007 | Jaspar<br>Transfac | *ftz-f1* |
| *ct* | 0.0151 | Jaspar | - |
| *D* | 0.0305 | Jaspar | *ftz-f1* |
| *opa* | 0.0002 | Jaspar | - |
| *Stat92E* | 0.0230 | Jaspar | - |
| *usp* | 0.0057 | Transfac | *ftz-f1*, *Smr* |

**b**

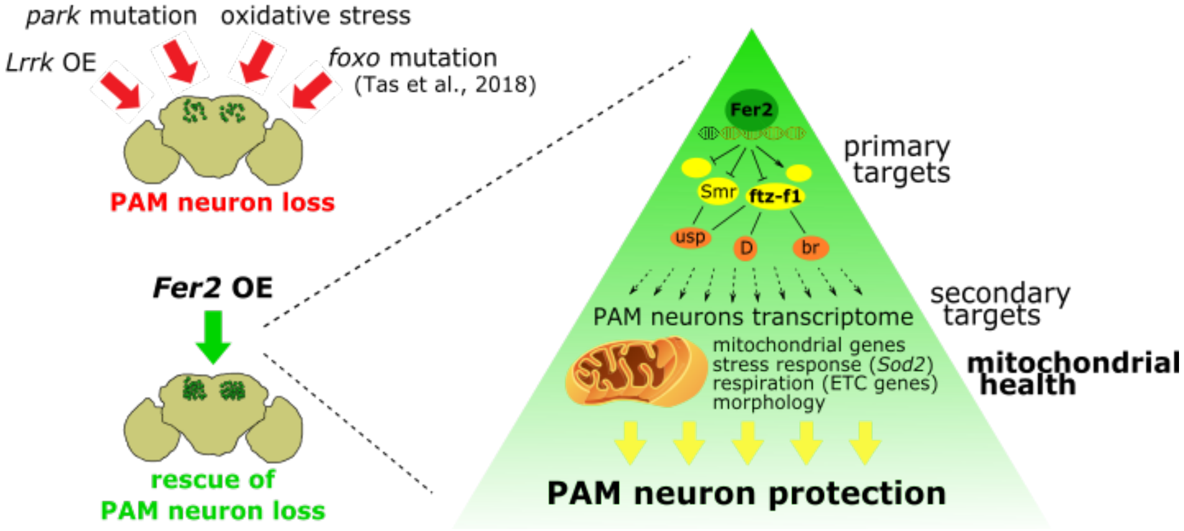


**Supplementary Figure 3. Genetic pathways downstream of *Fer2*.**
**(a)** List of transcription factors that are expressed in PAM neurons and whose binding sites are significantly overrepresented in the promoters of PAM RNA-seq DEGs. Enrichment p-value and the databases used for the analysis are indicated. Transcription factors known or predicted to interact with one or more *Fer2* direct targets are highlighted in orange. **(b)** Model of the role of *Fer2* in dopaminergic neuroprotection. *Fer2* binds to and regulates the expression of a set of direct target genes, including multiple transcription factors and chromatin regulators. *Fer2* direct targets FTZ-F1 and SMR interact with D, Br and USP and regulate a large set of genes within PAM neurons. In this manner, *Fer2* controls multiple pathways leading to regulation of mitochondrial gene expression, improved mitochondrial health, and PAM neuron survival against genetic and oxidative insults.

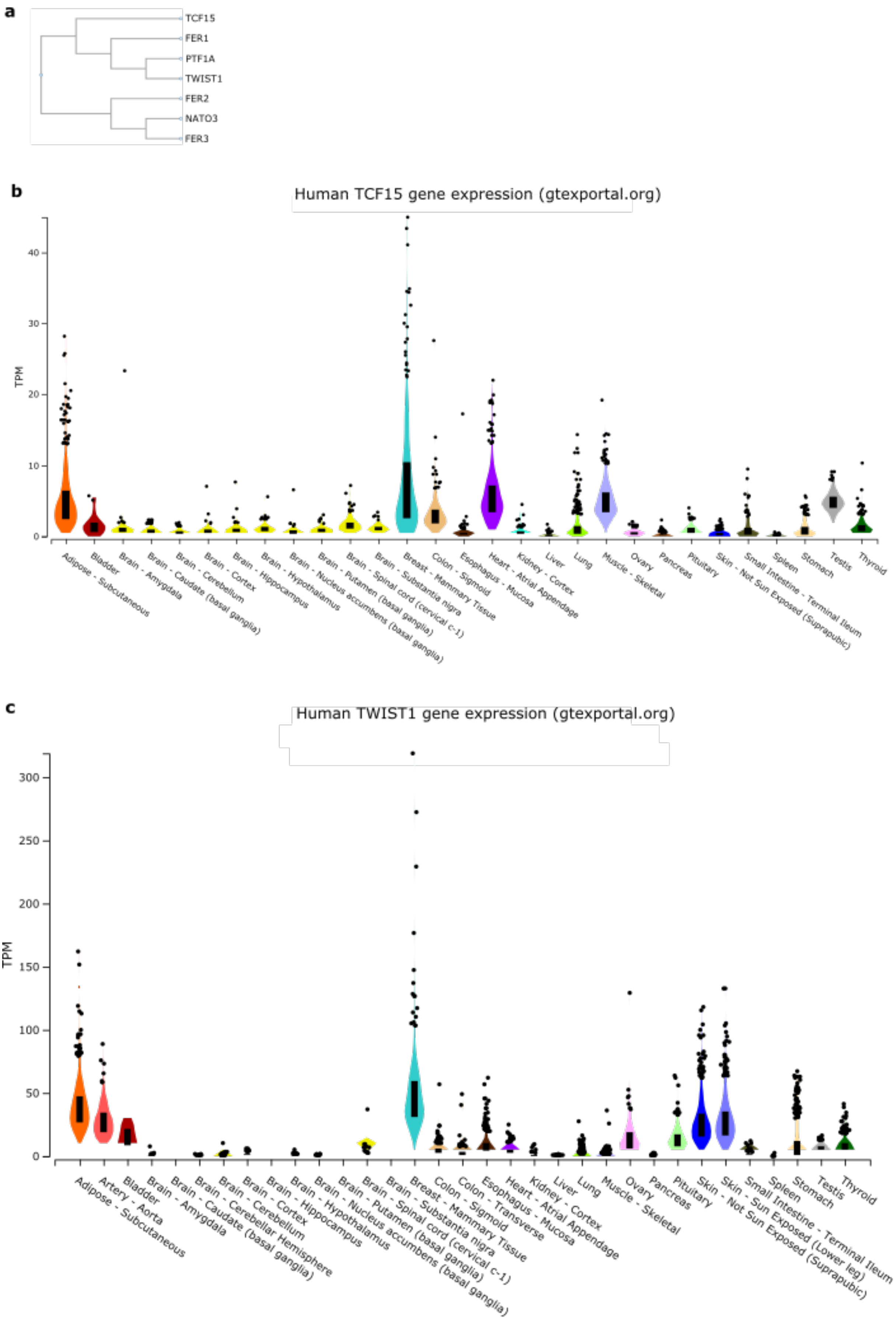
a
TCF15
FER1
PTF1A
TWIST1
FER2
NATO3
FER3
b
Human TCF15 gene expression (gtexportal.org)
TPM
0
10
20
30
40
Adipose - Subcutaneous
Bladder
Brain - Amygdala
Brain - Caudate (basal ganglia)
Brain - Cerebellum
Brain - Cortex
Brain - Hippocampus
Brain - Hypothalamus
Brain - Nucleus accumbens (basal ganglia)
Brain - Putamen (basal ganglia)
Brain - Spinal cord (cervical c-1)
Brain - Substantia nigra
Breast - Mammary Tissue
Colon - Sigmoid
Esophagus - Mucosa
Heart - Atrial Appendage
Kidney - Cortex
Liver
Lung
Muscle - Skeletal
Ovary
Pancreas
Pituitary
Skin - Not Sun Exposed (Suprapubic)
Small Intestine - Terminal Ileum
Spleen
Stomach
Testis
Thyroid
c
Human TWIST1 gene expression (gtexportal.org)
TPM
0
50
100
150
200
250
300
Adipose - Subcutaneous
Artery - Aorta
Bladder
Brain - Amygdala
Brain - Caudate (basal ganglia)
Brain - Cerebellar Hemisphere
Brain - Cerebellum
Brain - Cortex
Brain - Hippocampus
Brain - Hypothalamus
Brain - Nucleus accumbens (basal ganglia)
Brain - Putamen (basal ganglia)
Brain - Spinal cord (cervical c-1)
Brain - Substantia nigra
Breast - Mammary Tissue
Colon - Sigmoid
Colon - Transverse
Esophagus - Mucosa
Heart - Atrial Appendage
Kidney - Cortex
Liver
Lung
Muscle - Skeletal
Ovary
Pancreas
Pituitary
Skin - Sun Exposed (Lower leg)
Skin - Not Sun Exposed (Suprapubic)
Small Intestine - Terminal Ileum
Spleen
Stomach
Testis
Thyroid

d

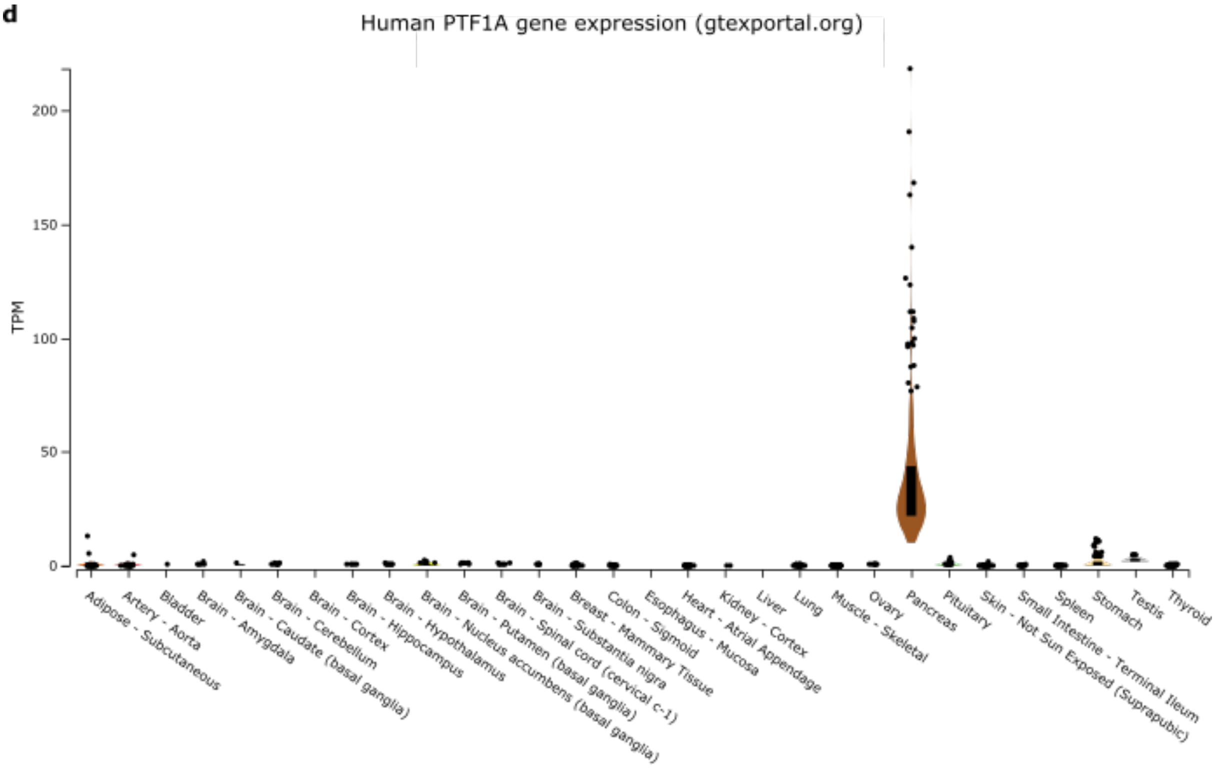


**Supplementary Figure 4. Phylogenetic and expression analysis of mammalian genes with significant similarity to FER2 protein sequence.**
**(a)** Phylogenetic tree generated by multiple protein sequence alignment of *Drosophila* FER1, FER2, FER2 and mouse TCF15, NATO3, PTF1A and TWIST1, using ClustalW. **(b-d)** *TCF15* (**b**), *TWIST1* (**c**) and *PTF1A* (**d**) expression levels in human tissues measured by RNA-seq as reported in GTEx portal (gtexportal.org).

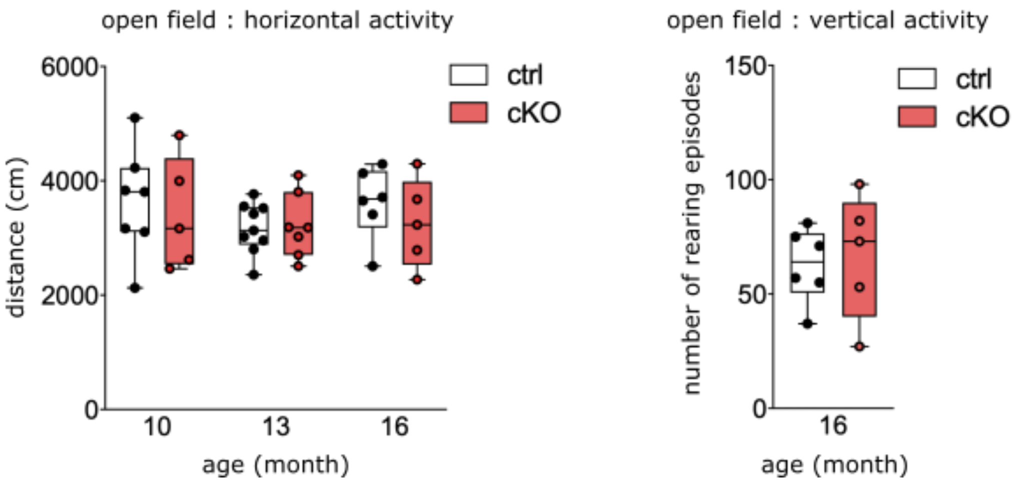


**Supplementary Figure 5. Locomotor activity measured in the open field test is not affected in *Nato3* cKO mice.**
Levels of horizontal activity (left), measured as the total distance traveled in the arena, and vertical activity (right), measured as the number of rearing events, were not different between Nato3 cKO and control mice. n=5-7 mice per group. No statistically significant difference between ctrl and cKO by Mann-Whitney test.
